# Molecular signatures of maladaptive plasticity in the amygdala in a rat model of chronic neuropathic pain

**DOI:** 10.64898/2026.01.05.697820

**Authors:** P. Presto, J. Cardenas, G. Ji, O. Ponomareva, V. Neugebauer, I. Ponomarev

## Abstract

Chronic pain, a complex multidimensional disorder, remains a major health care issue and a therapeutic challenge. Neuropathic pain is a chronic pain condition that results from damage or dysfunction in the nervous system. While mechanisms of neuropathic pain at the peripheral and spinal cord level have been extensively studied, pain mechanisms in the brain remain underexplored. The amygdala, a limbic brain region, has emerged as a critical brain area for the emotional-affective dimension of pain and pain modulation. Amygdala neuroplasticity has been associated with pain states, but exact molecular and cellular mechanisms underlying these states and the transition from acute to chronic pain are not well understood. Here, we used the spinal nerve ligation (SNL) model of neuropathic pain in male rats to investigate changes in gene expression in the amygdala at the chronic pain stage using RNA sequencing (RNA-Seq). Two amygdala nuclei, basolateral (BLA) and central (CeA), were investigated in a hemisphere-dependent manner. We used an integrative approach that focuses on functional significance and cell type specificity of differentially expressed genes (DEGs) to nominate mechanistic targets for central regulation of chronic pain. Our integrative transcriptomic and bioinformatic analyses identified individual genes (e.g., *Cxcl10, Cxcl12, Mbp, Plp1, Mag, Mog, Slc17a6, Gad1, Sst*), molecular pathways (e.g., cytokine-mediated signaling pathway), biological processes (e.g., myelination, synaptic transmission), and specific cell types (e.g., oligodendrocytes, glutamatergic and GABA-ergic neurons) affected by chronic pain. Our results also provide evidence for the emerging concept of hemispheric lateralization of pain processing in the amygdala. Overall, our study proposes oligodendrocyte dysfunction in the amygdala, neuroimmune signaling in CeA, and glutamatergic neurotransmission in BLA as mechanistic determinants of and potential therapeutic targets for the management of chronic neuropathic pain.

## Introduction

The International Association for the Study of Pain (IASP) defines pain as “an unpleasant sensory and emotional experience associated with, or resembling that associated with, actual or potential tissue damage” (Raja et al., 2020). Acute pain serves as a protective warning of potential danger and plays a critical role in the body’s healing response. In contrast, chronic pain is a disorder that impairs functioning and quality of life, necessitating effective treatment or management. Chronic pain such as neuropathic pain is difficult to treat, in part, due to its highly complex nature that involves brain processes that are still not well understood. Drug development and drug repurposing require understanding the mechanisms underlying maladaptive processes in the nervous system at every level associated with the transition from acute to chronic (chronification of) pain.

The amygdala, a limbic structure located in the medial temporal lobe, has emerged as a critical brain area for the emotional-affective dimension of pain and pain modulation based on preclinical (Apkarian et al., 2013; Neugebauer, 2015, 2020; Thompson & Neugebauer, 2017) and clinical (Kulkarni et al., 2007; Liu et al., 2010; Simons et al., 2014) evidence. The amygdala is comprised of anatomically and functionally distinct nuclei and subdivisions. The regions of the amygdala that are relevant for sensory and pain-related processing are the lateral/basolateral complex (LA/BLA), the intercalated cell (ITC) mass, and the central nucleus (CeA). The CeA is the major amygdala output center and contains efferent projections to the brainstem, hypothalamus, and basal forebrain regions (Becker & Carrasquillo, 2019; Neugebauer et al., 2020, 2023; Y. Wang et al., 2023). Changes in amygdala neuronal activity have been observed in pain models, and neuroplasticity within the amygdala has been linked to pain-like behaviors. Pain-related neuroplastic changes lead to hyperexcitability in amygdala output neurons (Neugebauer, 2020), driving pain-like behaviors in both acute (Kiritoshi & Neugebauer, 2018; Neugebauer & Li, 2003) and chronic (G. Ji et al., 2017; Jiang et al., 2014) pain models. Despite recent advances in dissecting brain neurocircuitry of pain-like behaviors, specific cellular and molecular mechanisms underlying the chronification of pain are not well understood.

Gene expression (transcriptomic) analysis has been widely used for gaining insight into molecular mechanisms of brain disorders and the discovery of novel molecular targets for medication development. Transcriptomic profiling in the periphery and spinal cord of animal pain models has revealed regulation of many genes implicated in several biological processes including neuronal functions and the immune response (Ray et al., 2019; Ray et al., 2023), though pain-related gene expression profiles within the brain are understudied. A recent study of the whole amygdala transcriptome at the acute stage (6 days) of a mouse model of neuropathic pain (Su et al., 2021) reported several individual genes and biological functional groups affected by the induction of pain at this stage. Important knowledge gaps, however, remain and these include the chronic pain condition, the role of specific cell types in different amygdala subregions, including non-neuronal cell types such as microglia and oligodendrocytes, and hemispheric lateralization. There is good evidence to suggest different roles of the right and left amygdala in the regulation of pain-like behaviors, and, potentially, chronification of pain (G. Ji & Neugebauer, 2009; Carrasquillo & Gereau, 2008; Allen et al., 2021). The molecular signatures of pain-related amygdala plasticity in the right and the left amygdala nuclei, which may drive pain-related behaviors remain to be determined.

Here, we characterized the transcriptional profiles of both the right and the left CeA and BLA of male rats at the chronic stage of neuropathic pain. We used the well-established spinal nerve ligation (SNL) animal model neuropathic pain, which mimics clinical symptoms like mechanical allodynia (pain from non-painful touch) and heat and cold hyperalgesia. We used published molecular markers of brain cell types to determine cellular identity of differentially expressed genes (DEGs) and to generate cell type-specific hypotheses of brain changes in neuropathic pain. Our data implicate neuroimmune signaling, myelination, and specific cell types in the amygdala in the chronification of neuropathic pain.

## Materials and methods

### Animals

Adult male Sprague-Dawley rats (250-400 g, 12 weeks of age at time of behavioral testing) were group-housed (n = 3 per cage) in a temperature-controlled room under a 12 hour day/night cycle with ad libitum access to food and water. All experimental procedures were approved by the Institutional Animal Care and Use Committee (IACUC, protocol #21026) of Texas Tech University Health Sciences Center (TTUHSC) and conformed to the guidelines of the International Association for the Study of Pain (IASP) and the National Institutes of Health (NIH).

### Neuropathic pain model

The well-established SNL rat model of neuropathic pain (Ho Kim & Mo Chung, 1992) was utilized to induce a stable and long-lasting peripheral neuropathy. Rats were anesthetized with isoflurane (2-3%; precision vaporizer, Harvard Apparatus, Holliston, MA) and underwent surgery to expose and tightly ligate the left L5 spinal nerve using 6-0 sterile silk. A sham-operated control group underwent a similar surgical procedure where the L5 spinal nerve was exposed but not ligated. Topical antibiotic (Bacitracin) was applied daily for 5 days after all surgical procedures to prevent infection.

### RNA isolation and bulk sequencing

At the chronic stage of neuropathic pain (4 weeks post-SNL or -sham surgery), rats (SNL, n = 6; sham, n = 6) were euthanized via decapitation. The brains were rapidly extracted and oxygenated in ice-cold ACSF that contained the following (in mM): 125.0 NaCl, 2.6 KCl, 2.5 NaH_2_PO_4_, 1.3 CaCl_2_, 0.9 MgCl_2_, 21.0 NaHCO_3_, and 3.5 glucose. Coronal brain slices (1,000 μM) containing the left and right CeA were prepared using a Vibratome (VT1200S, Leica Biosystems, Nussloch, Germany) as described previously (Hein et al., 2021; Presto & Neugebauer, 2022; Thompson et al., 2018). The left and right CeA and BLA were dissected from freshly harvested slices for bulk RNA sequencing analysis. Total RNA was isolated using the MagMAXTM-96 Kit (Life Technologies, Carlsbad, CA) and checked for quality control (all RIN values were > 8.4). RNA library preparation and sequencing were performed at the University of Texas at Austin Genomic Facility. Illumina Tag-Seq of polyA enriched total RNA sequencing was performed (single end, 100 bp).

### Quality control and alignment

Read quality was assessed using Fastp (v.0.23.4) (Chen et al., 2018). Reads with a Phred score below 30 were removed (-q 30), and we trimmed 10 low-quality nucleotides from the ends of the reads (-t 10), resulting in reads with a mean length of 91 bp. To align the reads to the reference genome, we utilized Salmon (v.1.10.2) (Patro et al., 2017). We created a decoy-aware index using the mRatBN7.2 genome release 111 from Ensembl (Cunningham et al., 2022). After building the index, we used the salmon quant command with the -l A flag to automatically identify the library type and the --validatingMappings flag to quantify transcript expression.

### Differential expression analysis

We imported the transcript expression data for each region (CeA, BLA) into RStudio using tximeta (v1.20.3) (Love et al., 2020) and used the summarizetogene function to create one edgeR object per region. Prior to differential gene expression analysis (DEG), we performed principal component analysis (PCA) using ggfortify (v0.4.17) to observe the main sources of variation and identify potential outliers. Then a DEG analysis was conducted using edgeR (v4.0.16) (Robinson et al., 2010). Prior to running the QLFtest, we applied the filterbyexpress function to remove low-expressed genes (defined as those with fewer than one counts per gene in 40% of the samples). We followed this with the calcnormfactors function to normalize the data. The makecontrast function was used to define the experimental group comparisons. Our experimental groups were: Sham.Left, Sham.Right, SNL.Left and SNL.Right. For each brain region, we identified genes with the main effect of Pain (SNL vs Sham), Side (Left vs Right) and their interaction. Genes with a nominal P-value < 0.05 were considered DEGs. False discovery rate (FDR) was also calculated. We visualized the results using volcano plots and Venn diagrams created with ggplot (v3.5.1), ggvenn (v0.1.10), and ggrepel (v0.9.5) (Wickham, 2009). To generate all heatmaps for DEGs and cell type -log(Pvalue), the package ComplexHeatmap (v2.22.0) was used, while all Pain x Side interaction plots for individual genes were created by ggplot (v3.5.1).

### Functional group over-representation analysis

We conducted functional group over-representation analysis for biological processes and molecular pathways using ClusterProfiler (v4.10.1), utilizing Gene Ontology (GO), Reactome, and the Kyoto Encyclopedia of Genes and Genomes (KEGG) databases (Kanehisa et al., 2023). Genes with a nominal p-value of less than 0.05 were used as queries, and all detected genes were used as background. Upregulated and downregulated genes were analyzed separately and together resulting in 3 lists of DEGs. Dot plots representing overrepresented terms were generated with ggplot2 (v3.5.1). To explore connections between overrepresented terms, we built gene networks using the Cytoscape (Shannon et al., 2003) (v3.10,3).

### Cell type over-representation analysis

To determine cell type-specific DEGs and cell types most responsive to the factors of Pain, Side and their interaction, we used published molecular markers of cell types in the amygdala and performed a cellular “deconvolution” and cell type over-representation analyses. We explored published single nucleus RNA sequencing data (Zhou et al., 2023; GEO accession GSE212417) to identify cell type-specific genes in the amygdala and used this information to define cellular identity of DEGs. We downloaded seven sample matrices from GSE212417 corresponding to the whole amygdala from treatment-naïve rats and used a published protocol (Marquez-Galera et al., 2022) and Seurat (v5.1.0) (Hao et al., 2021) to identify cellular clusters and define cell types by using canonical cellular markers. Thirteen cell types were defined based on the number of clusters from the dataset: mature oligodendrocytes (OD), immature oligodendrocytes (Immature OD), oligodendrocyte precursor cells (OPC), microglia, astrocytes, endothelial cells, VGUT1-positive glutamatergic neurons (ExNeuron), VGUT2-positive, Nos1-positive glutamatergic neurons (Nos1+), intercalated cells (IC Cells), general GABA-ergic inhibitory neurons (InhNeuron), proenkephalin-positive neurons (Penk+), somatostatin-positive neurons (Sst+), and reelin-positive neurons (Reln+). To identify amygdala cell types over-represented with DEGs, we ran a hypergeometric test (phyper) in R with a significance level of p<0.05 and an expression/enrichment filter of 0.6 for the genes from the referenced dataset for each region. Up- and down-regulated genes were analyzed separately and together, resulting in 3 lists of DEGs. To further investigate the effects of SNL on expression of *all cell type-specific genes* of five over-represented cell types (OD, Immature OD, OPC, ExNeuron, Nos1+) in CeA and BLA, we built distribution curves based on t-value statistic for each gene and compared mean t-values for each cell type to zero chance using a one sample t-test followed by Bonferroni correction. T-values were calculated from the edgeR-based F statistics (Supplemental Table 1).

## Results

### Effects of chronic pain (SNL model) on gene expression in BLA and CeA

We used the well-established SNL animal model of neuropathic pain to identify individual genes affected by chronic pain in CeA and BLA. PCA identified “brain region” as the main source of variability in gene expression, separating CeA and BLA samples along PC1 axis (Fig.1A), and providing the rationale for separate analyses within each brain region. Lists of DEGs showing main effect of Pain were largely different between BLA and CeA (Fig.1B). However, the inter-region overlap of 114 DEGs was significantly greater than expected by chance (hypergeometric p-value < 0.01), suggesting some common mechanisms of pain regulation in the two brain regions. These DEGs were further investigated using cell type over-representation analysis (presented below). Heatmaps in CeA (Fig.2A) and BLA (Fig.3A) show selected DEGs up- or down-regulated in the SNL group compared to control. Overall results can be found in Supplemental Table 1.

**Figure 1.**
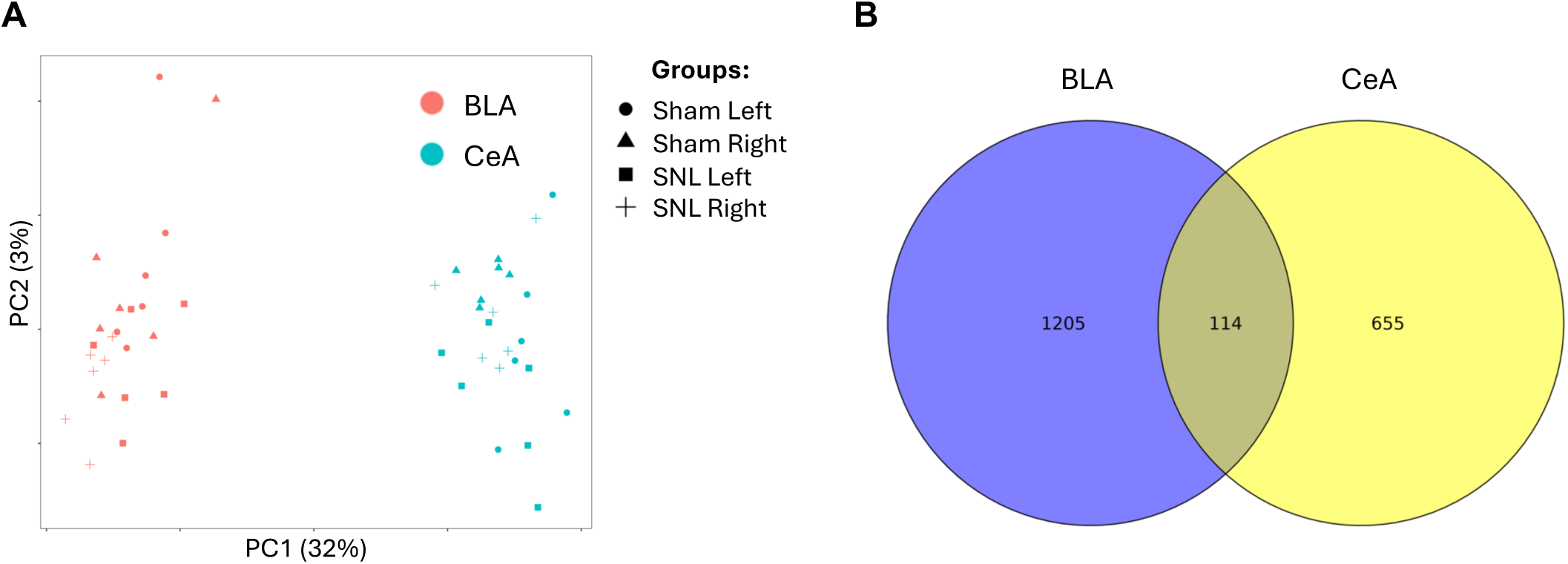
Principal Component Analysis (PCA) and differentially expressed genes (DEGs). A. PCA shows that gene expression in BLA and CeA are mainly different. PCA shows that the main source of variability in brain gene expression is ”brain region”, as the Principal Component 1 (PC1), which accounts for the largest portion of variance in gene expression (32%), separates BLA and CeA into distinct clusters. This is consistent with the literature. Therefore, we analyzed BLA and CeA separately for the effects of SNL vs Sham. B. BLA and CeA showed largely different lists of DEGs, however the overlap of 114 DEGs was significantly greater than expected by chance (hypergeometric p-value < 0.01), suggesting some common mechanisms. Further analysis showed that the list of overlapping genes was over-represented with molecular markers of oligodendrocytes, with most of them being down-regulated in the SNL group.

**Figure 2.**
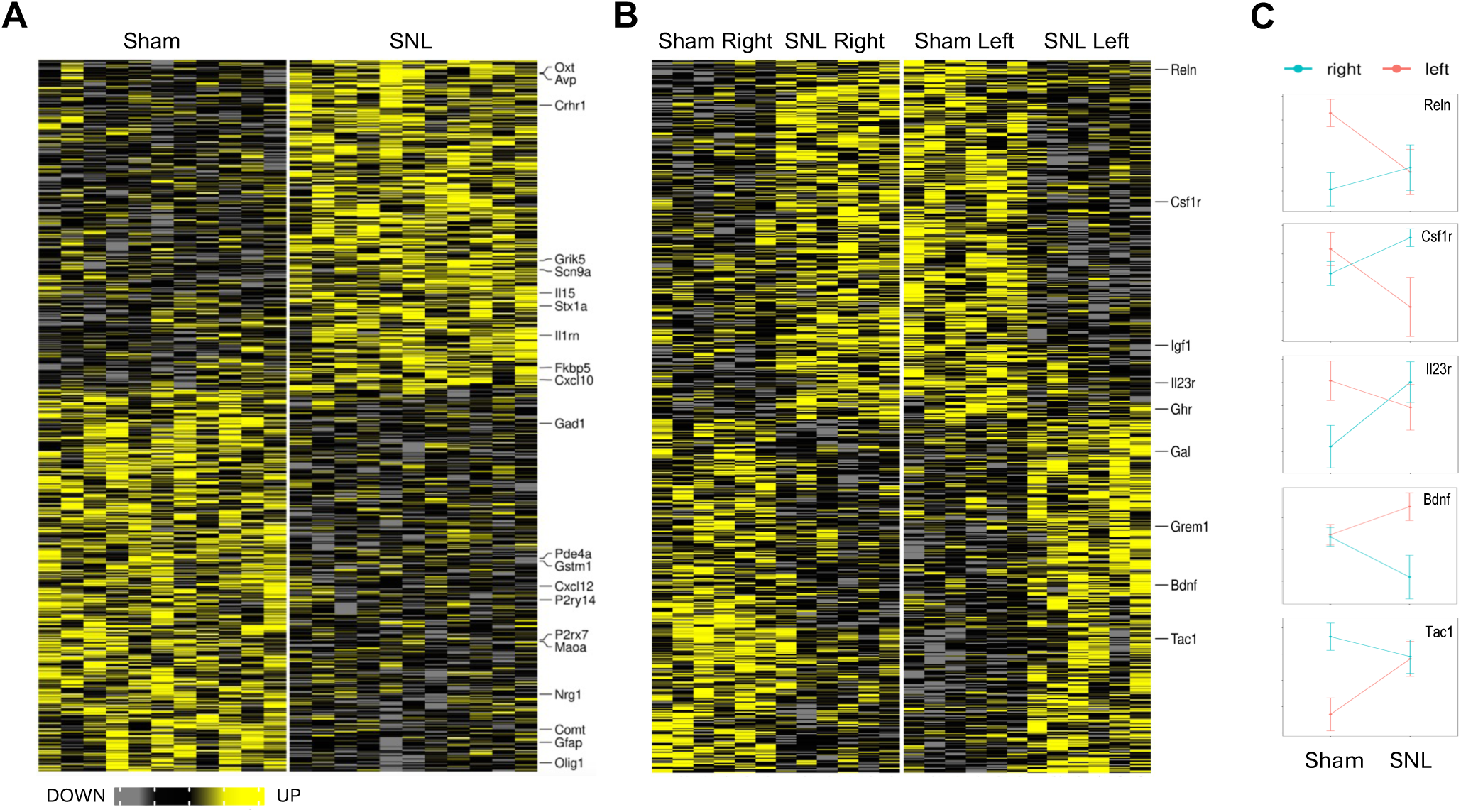
Differentially expressed genes (DEGs) in CeA. Heatmap plots of individual samples (columns) show DEGs for the main effect of Pain (A) and Pain by Side interaction (B). Gene symbols of selected genes with known or proposed function in pain modulation are shown to the right of each heatmap. Genes with statistically significant Pain x Side interaction are affected differently by SNL in the left vs right hemisphere. Normalized Means ± SEM for selected genes with the interaction effect are shown in C.

**Figure 3.**
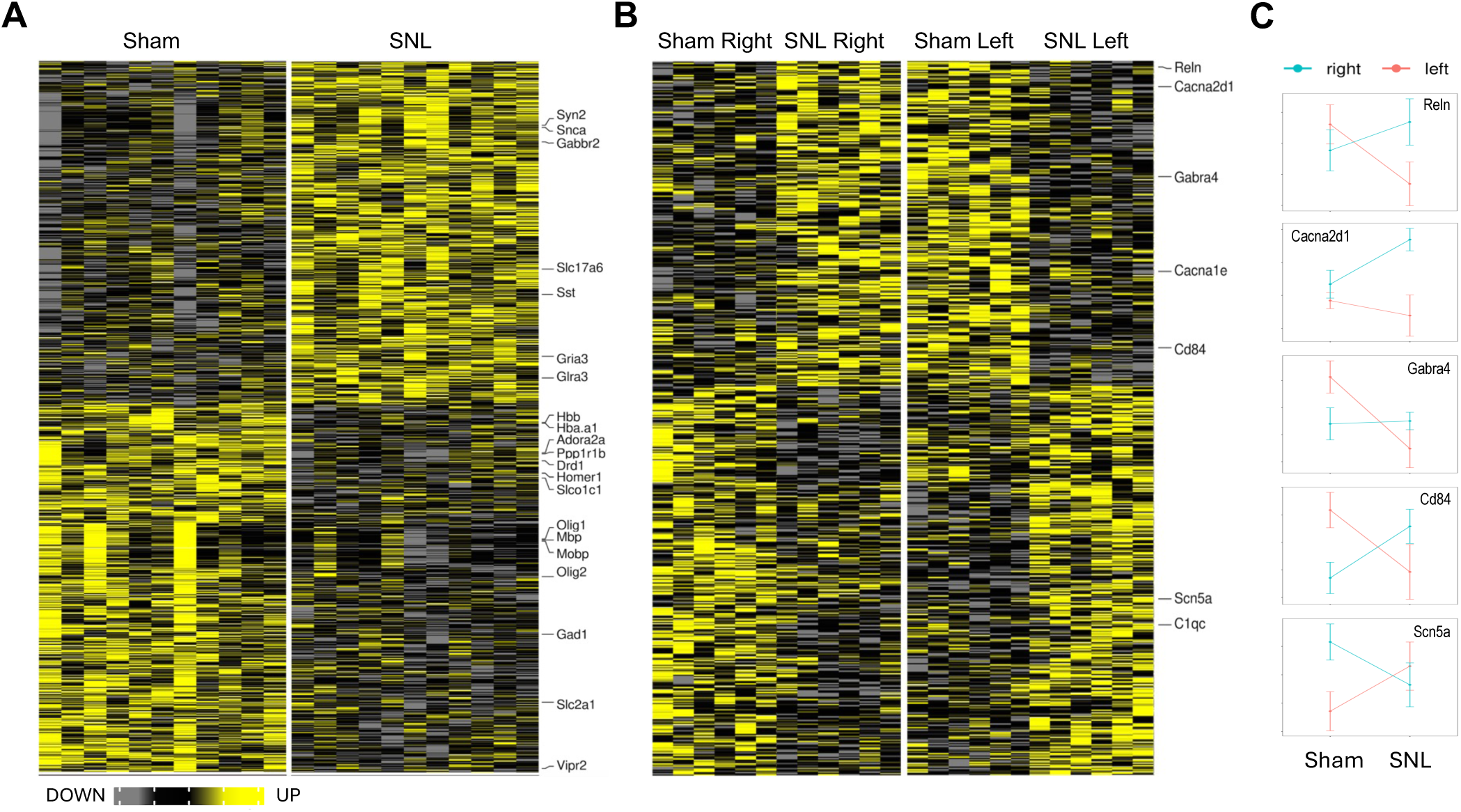
Differentially expressed genes (DEGs) in BLA. Heatmap plots of individual samples (columns) show DEGs for the main effect of Pain (A) and Pain by Side interaction (B). Gene symbols of selected genes with known or proposed function in pain modulation are shown to the right of each heatmap. Genes with statistically significant Pain x Side interaction are affected differently by SNL in the left vs right hemisphere. Normalized Means ± SEM for selected genes with the interaction effect are shown in C.

In CeA, we identified several DEGs implicated in synaptic transmission (e.g., *Stx1a, Crhr1, Oxt*) and neuroimmune signaling (e.g., *Il1rn, Cxcl10, Cxcl12, P2ry14*), suggesting a potential interplay of these processes in pain regulation. In BLA, several DEGs were also implicated in synaptic transmission (e.g., *Gria3, Drd1, Homer1*), with some genes being molecular markers of specific neuronal types (e.g., *Slc17a6* in glutamatergic neurons, *Gad1* and *Sst* in GABAergic neurons). In addition, several molecular markers of oligodendrocytes implicated in myelination (e.g., *Mbp, Plp1, Myrf, Olig1*) were down-regulated in the SNL group, suggesting BLA hypomyelination in chronic pain. *Myrf* and *Olig1* were also down-regulated in CeA.

### Hemispheric lateralization in pain-induced gene expression

Accumulating evidence suggests right-hemispheric lateralization of pain processing in the amygdala, though relatively little is known about the role of lateralization in chronic pain (G. Ji & Neugebauer, 2009; Carrasquillo & Gereau, 2008; Allen et al., 2021). To investigate the potential role of hemispheric lateralization in pain-induced effects on gene expression, we identified DEGs showing Pain x Side interaction. The majority of such genes showed side-specific regulation by chronic pain, i.e., regulated in the opposite directions in the right and left hemispheres 4 weeks after the SNL surgery. Overall results of this analysis are shown in Figures 2BC and 3BC for CeA and BLA respectively as well as in Supplemental Table 1.

In CeA, we identified several neuronal (e.g., *Reln, Bdnf, Tac1*) and neuroimmune (e.g., *Csf1r, Il23r*) genes with the interaction effects, which was consistent with the main effect findings. For example, brain-derived neurotrophic factor (*Bdnf*) gene was down-regulated in the right, while up-regulated in the left hemisphere. In BLA, the majority of DEGs were neuronal (*Reln, Gabra4, Scn5a*). Figures 2C and 3C show directionality of SNL-induced changes in the right and left hemispheres.

### Functional group over-representation analysis

We further investigated the potential role of different biological categories and molecular pathways by performing functional group over-representation analysis for DEG lists. Representative results of this analysis for the effect of Pain (SNL vs Sham) in CeA and BLA are shown in Figure 4, and all results can be found in Supplemental Table 2. Over-represented functional groups (OFG) common for CeA and BLA included myelination and oligodendrocyte development, both mainly containing down-regulated genes, while CeA-specific OFGs tended to include more immune-related categories, such as TNF signaling pathway and cytokine-mediated signaling pathway, and BLA-specific OFGs tended to include more neuron-related categories, such as glutamatergic synapse and regulation of synaptic plasticity.

**Figure 4.**
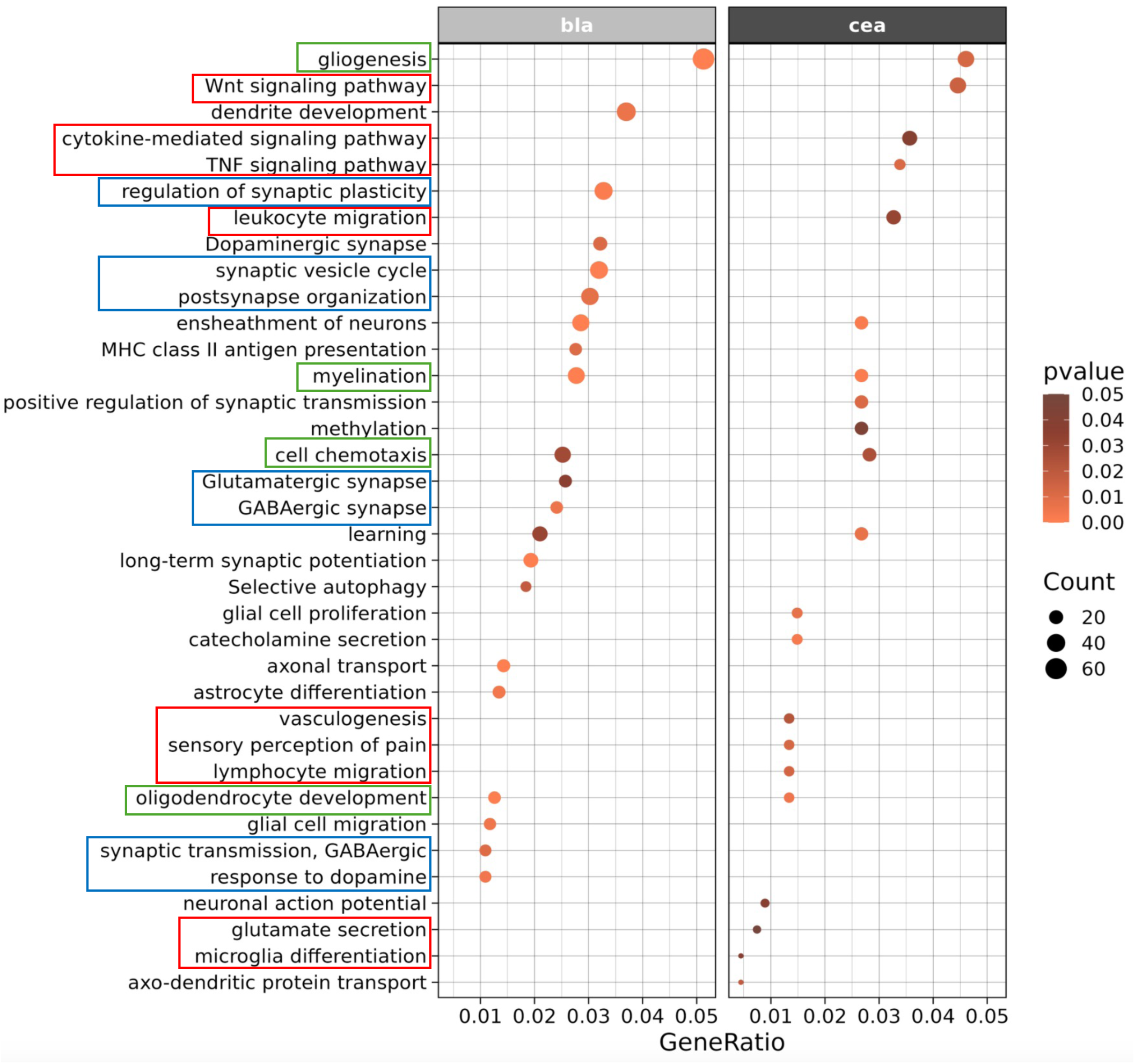
Representative results of functional group over-representation analysis for the effect of Pain (SNL vs Sham) in BLA and CeA. Green boxes outline selected functional categories similarly affected in both brain regions, while red (CeA) and blue (BLA) boxes outline brain region – specific functional groups.

To identify DEGs common for immune-, glia- and neuron-related processes in CeA, we compared gene lists of these categories and determined several genes linking them, including *Cxcl10, Cxcl12, Il1rn,* and *Cd200* (Fig.5). For example, *Cxcl10* was up-, while *Cxcl12* was down-regulated in SNL animals. These genes are chemokines shown to be involved in the communication between neurons and glia, thus, implicating the interplay between these cell types in regulation of neuroimmune signaling in chronic pain.

**Figure 5.**
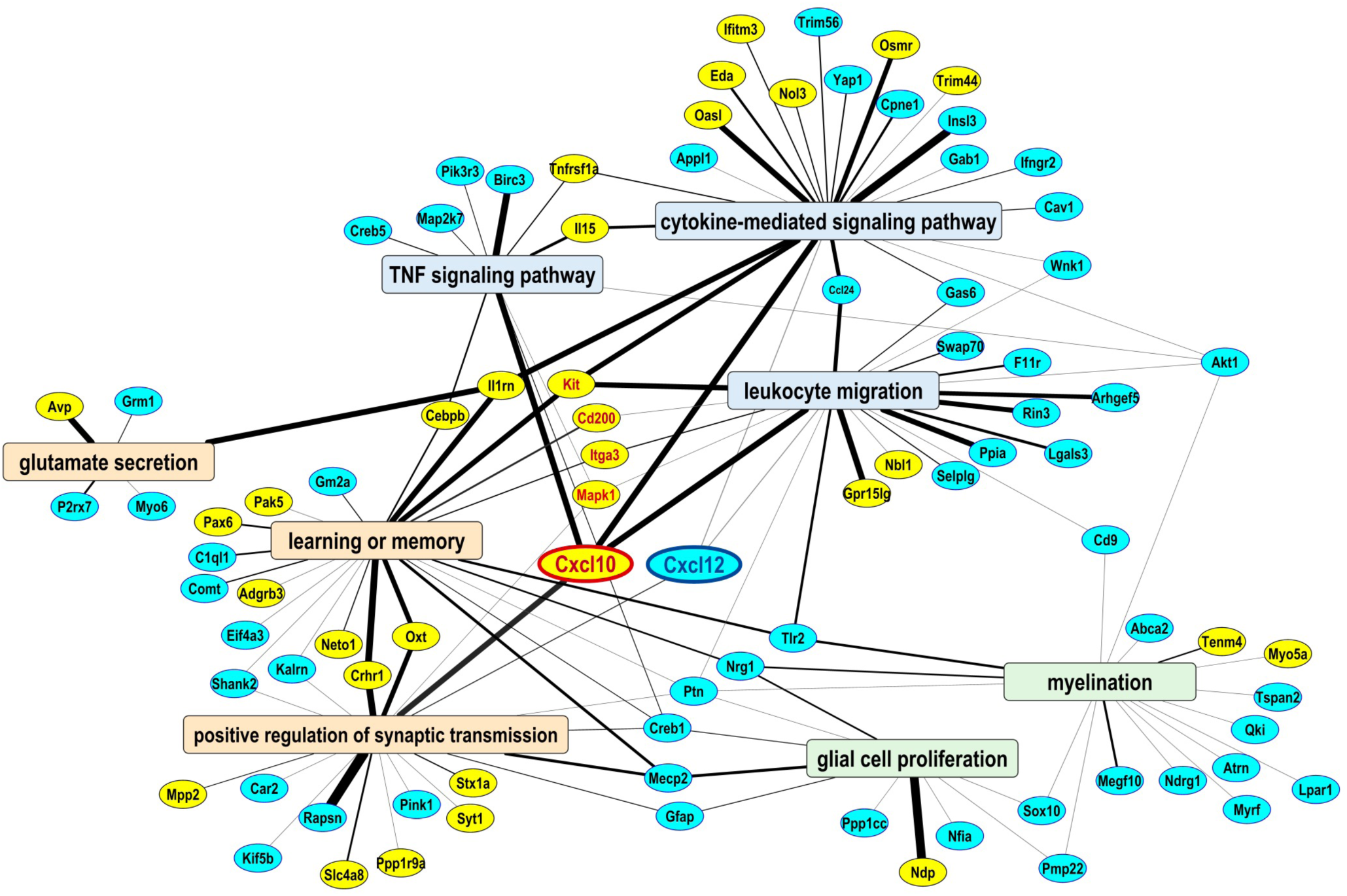
Representative network of over-represented functional groups and associated DEGs in CeA implicated in the regulation of neuropathic pain. Functional groups from three “domains” are shown: “neuronal” (orange background), “immune” (blue background), and “glial” (green background). DEGs (yellow = up-regulated, blue = down-regulated in SNL vs Sham group) connecting these domains are potential CeA targets for cell-cell interactions in the context of neuropathic pain. Highlighted are 2 genes in the middle, Cxcl10 and Cxcl12 chemokines implicated in communication between microglia, astrocytes and neurons.

### Cell type over-representation analysis

To explore the potential role of different cell types in the amygdala in pain regulation we identified cell type-specific DEGs using published datasets and performed cell type over-representation analysis. This analysis revealed two main findings: 1) down-regulated genes in CeA and BLA were over-represented in non-neuronal cell types, especially in ODs and immature ODs, and 2) up-regulated genes in BLA were over-represented in neurons, especially glutamatergic neurons, both VGLUT1-positive ExNeuron and VGLUT2-positive Nos1+ (Fig. 6A). Distributions of t-statistics for all cell type-specific genes expressed in OD, immature OD, OPC, ExNeuron, and Nos1+ cell types validated these findings, showing overall shifts to the left (down-regulation) or to the right (up-regulation) from zero. Examples of cell type-specific DEGs include OD-specific and down-regulated *Olig1* and *Myrf* in both CeA and BLA, ExNeuron-specific and up-regulated *Gabbr2* and *Gria3*, and Nos1+-specific and up-regulated *Slc17a6* (VGLUT2) and *Gabrg2* in BLA. In addition, all OD-specific major myelin-associated genes (*Mbp, Plp1, Mag, Mog, Mobp*) were down-regulated in BLA, suggesting a loss of myelin in this region in the SNL group. Consistent with functional group and cell type over-representation analyses, the list of 114 DEGs commonly regulated between CeA and BLA (Fig.1B) was over-represented with OD-specific genes, all of them down-regulated (hypergeometric p-value<0.01), suggesting OD dysfunction in both regions as a common mechanism of neuroplastic changes in the amygdala in chronic pain.

**Figure 6.**
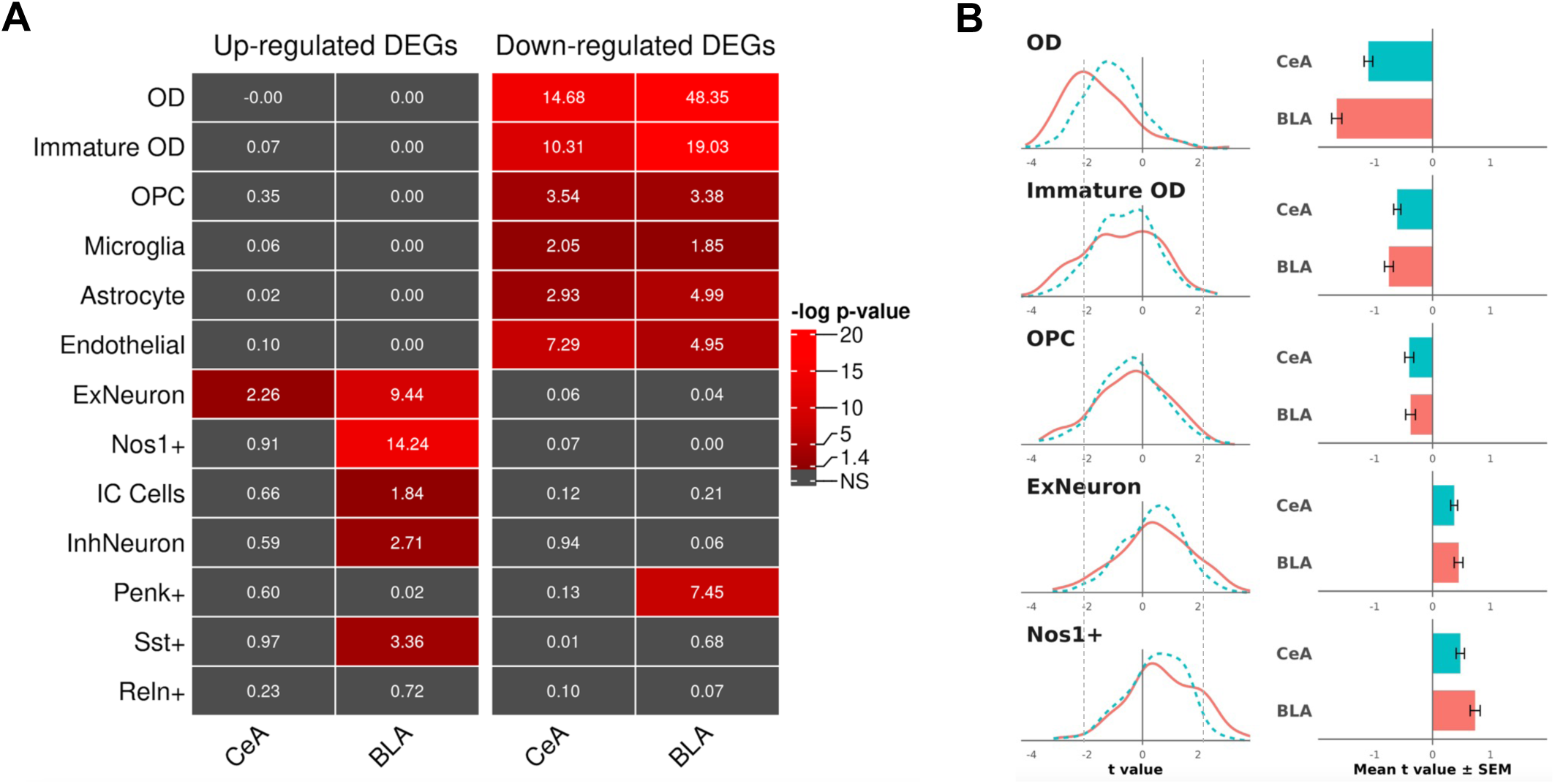
Cell type analysis of genes for the Pain condition (SNL vs Sham) in BLA and CeA. We used publicly available cell type-specific molecular markers to predict cellular identity of all genes with detectable expression in the amygdala. A. Cell type over-representation analysis of up- and down-regulated DEGs. Shown p-values are results of hypergeometric tests comparing the numbers of cell type-specific DEGs to chance. For example, DEGs with cell type-specific expression in oligodendrocytes (OD), astrocytes, and endothelial cells were mainly down-regulated in SNL rats in both CeA and BLA, while many neuron-specific DEGs were up-regulated in SNL animals in BLA. B. Distributions of t-statistics of all genes enriched in one of five cell types: mature ODs, immature ODs, OPCs, VGLUT1-positive glutamatergic neurons (ExNeuron), and VGLUT2- and Nos1-positive glutamatergic neurons. A distribution shift to the left suggests a general down-regulation of cell type-specific genes in SNL vs Sham, while a distribution shift to the right suggests a general up-regulation. Mean t-values for all five cell types (right panels) were significantly different from zero (one sample t test with Bonferroni correction, all p-values < 0.01). Dashed lines next to the -2 and +2 t values indicate thresholds for statistical significance for individual genes. For example, genes with t values < -2 or > +2 are considered DEGs.

## Discussion

In this study, we targeted two regions of the amygdala, CeA and BLA, for analysis of neuroplastic changes in chronic pain at the gene expression level, using transcriptomic and bioinformatic approaches, and identified individual genes, molecular pathways, biological processes, and specific cell types affected by chronic neuropathic pain in male rats. We used an integrative approach that focuses on functional significance and cell type specificity of DEGs to nominate mechanistic targets for central regulation of chronic pain. Our results are consistent with the literature and propose novel avenues for pain research. The analysis of gene expression changes in a chronic pain condition in specific cell types in different amygdala subregions and hemispheric lateralization using cellular “deconvolution” of transcriptomic data to generate novel cell type-specific hypotheses significantly advances findings from a previous study at the acute stage of the SNL model (Su et al., 2021).

We first compared SNL-induced transcriptomes between the two brain regions and found that there was little overlap between DEGs in the CeA and BLA. This finding was not surprising given that the two regions have distinct afferent and efferent projections and functional properties (Becker & Carrasquillo, 2019; J. Kim et al., 2017; McCullough et al., 2018; Neugebauer et al., 2023; Sah et al., 2003). The BLA is predominately composed of pyramidal glutamatergic projection neurons that receive polymodal sensory and nociceptive information from thalamic and cortical areas, including the anterior cingulate cortex (ACC), insula, and medial prefrontal cortex (mPFC) (Janak & Tye, 2015; Neugebauer, 2015; Neugebauer et al., 2004; Veinante et al., 2013b). The BLA integrates emotional-affective information and then relays this into the CeA for further processing (Thompson & Neugebauer, 2017). The CeA serves as the interface between nociceptive and affective processing and as a major output nucleus that connects to other brain regions involved in behavioral regulation. The BLA-CeA circuit has been implicated in the generation and modulation of pain-like behaviors (Corder et al., 2019; Neugebauer, 2020; Thompson & Neugebauer, 2017). Despite largely different effects of Pain on CeA and BLA transcriptomes, the overlap between DEGs in these regions was significantly greater than expected by chance. Many DEGs in this list were highly enriched in oligodendrocytes, all of them down-regulated, suggesting OD dysfunction in chronic pain. Oligodendrocytes support neurons by creating the myelin sheath for rapid signal transmission, providing metabolic support by supplying energy metabolites, and contributing to immune regulation. These functions are crucial for efficient neuronal communication, maintaining the integrity of axons, and supporting the overall health of the central nervous system (CNS).

Our comprehensive cell type-specific analysis found a drastic pain-induced decrease in expression of OD-specific genes in the amygdala, BLA to a greater extent. Many of OD-specific DEGs are implicated in the regulation of myelination, suggesting that chronic pain results in general OD dysfunction and demyelination. These findings are somewhat consistent with results from a recent study that explored neural circuits involved in comorbid chronic pain and depression (Becker et al., 2023). They found that hyperactivity of the neuronal pathway linking the BLA to the ACC was critical for chronic pain-induced depression, and activation of this pathway in control male mice was sufficient to trigger depressive-like behaviors. Furthermore, transcriptomic analysis revealed that BLA-ACC hyperactivity was associated with an enrichment for genes expressed by oligodendrocytes in the ACC; the majority of these were downregulated and were associated with decreased myelination pathways in this region. Given the central role of oligodendrocytes in maintaining axonal integrity and metabolic support, such reductions are expected to have profound functional consequences for local and long-range circuits.

Another main finding in our study is the potential role of neuroimmune signaling in CeA. Neuroimmune signaling is now recognized as an important peripheral and spinal pain mechanism (Grace et al., 2021), but the role of neuroimmune factors in pain-related amygdala neuroplasticity and behavior is not well understood, and little is known about the role, regulation and therapeutic potential of molecular crosstalk between neuronal and glial cell types in the brain in the context of pain. Our results suggest a potential interplay between neuronal and immune processes in pain regulation and nominate several molecular targets as mechanistic candidates for communication between neurons, microglia and other non-neuronal cell types. For example, we identified two chemokines, *Cxcl10* and *Clcx12*, up- and down-regulated respectively in the SNL group, which connected the “immune” and “synaptic” domains in CeA (Fig.5).

CXCL10 expression is generally low in the healthy CNS but is significantly up-regulated during inflammation, infection, and injury (Koper et al., 2018). Microglia and astrocytes are major sources of CXCL10, especially under inflammatory conditions (Michlmayr, D., & McKimmie, C.S., 2014). CXCL10 primarily acts via its receptor, CXCR3, which is expressed on neurons and can modulate synaptic transmission and promote apoptosis of neurons in a dose-dependent manner during chronic inflammation (Qiao et al., 2022). CXCL10 may be involved in the late phase of neuropathic pain and is up-regulated in astrocytes of the spinal cord after nerve injury or ischemia (Lu & Gao, 2023). CXCL12 is constitutively and widely expressed in the healthy CNS and is crucial for maintaining brain homeostasis (Li & Ransohoff, 2008). CXCL12 primarily signals through its receptors CXCR4 and, in the adult brain, it modulates neurotransmission, enhances synaptic plasticity, and generally provides neuroprotective effects against various insults (Wojcieszak et al., 2022). Interestingly, CXCL12 protein was up-regulated in dorsal root ganglia and spinal cord one to fourteen days after spared nerve injury in rats (a neuropathic pain model), but returned to the baseline level by day 21 (Bai et al., 2016). A down-regulation of *Cxcl12* in our study may indicate a different role of this gene at the chronic stage of neuropathic pain, CeA vs spinal cord differences, a potential mRNA expression compensation for the up-regulated protein, or all of the above. Taken together, our data suggest a proinflammatory state and disrupted homeostasis in CeA under chronic pain conditions.

We also investigated the potential role of hemispheric lateralization in pain-induced effects on gene expression by identifying DEGs showing Pain x Side interaction. The premise is based on the hypothesis that some genes may be regulated in the opposite directions in the right and left hemispheres under pain conditions. Similar to the main effect of Pain, many DEGs with the interaction effect in CeA were implicated in either neuronal or immune functions (e.g., *Tac1, Bdnf, Il23r)*. Tachykinin (Tac1) and brain-derived neurotrophic factor (BDNF) are known targets in pain research (Moy et al., 2019). For example, Substance P, the peptide encoded by the Tac1 gene, acts as an excitatory neurotransmitter in pain signaling pathways and is widely expressed in pain-related circuitry, including the spino-parabrachio-amygdala pain pathway that provides nociceptive information to the CeA (Jasmin et al., 1997, Barik et al., 2021, Torres-Rodriguez et al., 2024). Also, blocking the signaling of spinal BDNF and its TrkB receptor reversed mechanical allodynia induced by peripheral nerve injury in male rats (Coull et al., 2005). Both *Tac1* and *Bdnf* had at least a tendency for pain-induced down-regulation in the right and an up-regulation in the left hemispheres, consistent with structural synaptic changes of parabrachial input to CeA in chronic pain (Gandhi et al., 2021). DEGs with the interaction effects in BLA were mainly associated with neuronal functions. For example, reelin (*Reln*), a large extracellular matrix protein that plays an important role in brain development and function (Jossin, 2020), was down-regulated in the left and had a tendency to be up-regulated in the right hemisphere in SNL rats. In the adult brain, reelin is mainly secreted by a subset of GABA-ergic interneurons including the amygdala (Alexander et al., 2023), however, the specific role of these neurons in pain conditions has not been directly characterized. Taken together, our results suggest that hemispheric effects are important in the regulation of chronic pain.

Our cell type over-representation analysis showed that up-regulated DEGs in BLA were highly over-represented in glutamatergic neurons (both VGLUT1-expressing and VGLUT2-expressing Nos1-positive neurons). Many of these DEGs are implicated in the regulation of neuronal excitability and synaptic functions (e.g., VGLUT2), and this up-regulation may indicate an increased excitability of BLA glutamatergic neurons. Nos1-positive neurons are a distinct subpopulation of excitatory neurons that may be involved in pain-related anxiety and depression. The BLA Nos1-positive and Nos1-negative neurons have opposite effects on anxiety and depression-like behaviors by projecting to different areas (Cai et al., 2023). Together with the evidence for demyelination in this brain region, our results indicate an imbalance in neurotransmission and point to BLA as an important player in the regulation of chronic neuropathic pain.

In summary, our study provides novel insights into central regulation of chronic neuropathic pain centered on neuroimmune signaling and myelin changes in the amygdala. Although limited to a later stage of chronic pain and males, this research paves the way for future studies examining mechanisms of transition from acute to chronic pain, sex differences, the role of individual cell types and cell-cell interactions in the right and left amygdala as well as amygdala-related neurocircuits in chronification of pain. Overall, our data point to oligodendrocyte dysfunction in CeA and BLA, neuroimmune signaling in CeA, and glutamatergic neurotransmission in BLA as chronic pain mechanisms and potential therapeutic targets for the management of chronic neuropathic pain.

## Acknowledgements

We would like to thank Christian Bustamante, Brent Kisby, and Khadijah Mazhar for their help with initial bioinformatics analysis. This work was funded by NIH (R01 NS038261 to VN and IP and R01 AA027096 to IP).

